# Reproductive Compatibility in *Capsicum* is not Reflected in Genetic or Phenotypic Similarity Between Species Complexes

**DOI:** 10.1101/2020.11.30.403691

**Authors:** Catherine Parry, Yen-wei Wang, Shih-wen Lin, Derek W. Barchenger

## Abstract

Wild relatives of domesticated *Capsicum* represent substantial genetic diversity and thus sources of traits of potential interest. Furthermore, the hybridization compatibility between members of *Capsicum* species complexes remains unresolved. Improving our understanding of the relationship between *Capsicum* species relatedness and their ability to form hybrids is a highly pertinent issue. Through the development of novel interspecific hybrids in this study, we demonstrate interspecies compatibility is not necessarily reflected in relatedness according to established *Capsicum* genepool complexes. Based on a phylogeny constructed by genotyping using single sequence repeat (SSR) markers and with a portion of the *waxy* locus, and through principal component analysis (PCA) of phenotypic data, we clarify the relationships among wild and domesticated *Capsicum* species. Together, the phylogeny and hybridization studies provide evidence for the misidentification of a number of species from the World Vegetable Center genebank included in this study. The World Vegetable Center holds the largest collection of *Capsicum* genetic material globally, therefore this may reflect a wider issue in the misidentification of *Capsicum* wild relatives. The findings presented here provide insight into an apparent disconnect between compatibility and relatedness in the *Capsicum* genus, which will be valuable in identifying candidates for future breeding programs.

---

The genus *Capsicum* is comprised of about 35 species including five domesticated species: *C. annuum* (L.), *C. baccatum* (L.), *C. chinense* (Jacq.), *C. frutescens* (L.), and *C. pubescens* (Ruiz & Pav.) (Khoury et al., 2020). The diversity of *Capsicum* species represents a valuable genetic resource for crop improvement (Barchenger and Bosland, 2019). The primary limitations to improving productivity and quality of *Capsicum* are abiotic and biotic stresses, many of which lack sources of host tolerance or resistance (Barchenger et al., 2019). Furthermore, as a widely consumed crop with cultural and culinary value across global cuisines, there is high demand for *Capsicum* (Bosland and Votava, 2012). There is therefore significant incentive to overcome challenges to cultivation, and one means of doing so being the introgression of resistance to the various stresses that limit production of *Capsicum* species.

Understanding interspecies compatibility and identifying barriers to hybridization is essential to the design of introgression breeding programs. *Capsicum* species are divided among 11 clades (Bosland and Votava, 2012; Carrizo García et al., 2016) and grouped into three complexes - Annuum, Baccatum and Pubescens - based on their relative reproductive compatibility (Tong and Bosland, 1999; Pickersgill, 1971; Emboden Jr., 1962). There is understood to be relatively low reproductive compatibility between species complexes (van Zonneveld et al., 2015). However, a number of cross-complex hybridizations have been achieved (Eggink et al., 2014; Costa et al., 2009; Kamvorn et al., 2014; Yoon et al., 2006; OECD, 2006; Pickersgill, 1991), and the pre- and post-zygotic barriers to hybridization between genetic complexes are not fully understood (Yoon et al., 2006). This suggests isolation between complexes is not total, and there is therefore potential for introgression breeding, or design of genetic bridge strategies in order to best exploit this genetic variation.

In contrast to other Solanaceae crops, including tomato (*Solanum lycopersicum*) (Lin et al., 2014), potato (*S. tuberosum*) (Hirsch et al., 2013) and to a lesser extent eggplant (*S. melongena*) (Gramazio et al., 2017) introgression breeding using wild species has been relatively underutilized in *Capsicum*; (Mongkolporn and Taylor, 2011). The wild progenitor, *C. annuum* var. *glabriusculum* is a potential source of disease resistance, with reported resistance to Beet curly top virus (BCTV: *Curtovirus*) (Bosland, 2000; Jimenez, 2019). Members of the wild species *C. chacoense* and *C. rhomboideum* have been identified as being resistant to powdery mildew (*Leveillula taurica*) (McCoy and Bosland, 2019). Recently, an accession of *C. galapagoense* has been proposed to be a potential source of resistance to the insect pest, whitefly, based on trichome density and type (M. Rhaka, pers. comm.). However, despite extensive hybridization no successful progeny have so far been developed (Lin et al., 2020). These results are surprising, because *C. galapagoense* has been reported as part of the *C. annuum* clade, and readily hybridize with *C. annuum* accessions (Carrizo García et al., 2016; Pickersgill, 1971). One reason for unsuccessful hybridization attempts may be misidentification; several genebanks have incorrectly reported accessions identified as *C. galapagoense* which are, in fact, *C. frutescens* (P.W. Bosland, pers. comm.). Such misidentification presents a challenge to utilizing knowledge of the relatedness of *Capsicum* species and their ability to hybridize. Although the genetic diversity and variation within wild populations of *Capsicum* has been studied (Carrizo García et al., 2016; Cheng et al., 2016; Aguilar Meléndez et al., 2009; Oyama et al., 2006; Votava et al., 2002; Loaiza-Figueroa et al., 1989), the pool of phenotypic data for wild *Capsicum* species remains limited (Barchenger and Bosland, 2019). There also remains a lack of access to publicly available germplasm representing the diversity of wild *Capsicum* (Khoury et al., 2020). There is therefore an immediate need to better understand the role of wild *Capsicum* species in future breeding programs.

The objectives of this study were to elucidate the relationship between interspecies compatibility and relatedness through extensive interspecific hybridization and the construction of a phylogeny. We aimed to clarify the relationships among the wild and domesticated *Capsicum* species included in the study, and confirm the identities of several World Vegetable Center genebank accessions.

## Materials and Methods

Thirty-eight accessions of 15 species of *Capsicum* were chosen for this experiment (Table 1). The accessions were provided to the World Vegetable Center, having been collected from diverse locations and deposited into collections at either the World Vegetable Center Genebank, the World Vegetable Center Pepper Breeding Collection in Tainan, Taiwan, the United States Department of Agriculture - Agriculture Research Service National Plant Germplasm System, or the Chile Pepper Institute, New Mexico State University, Las Cruces, NM USA. Of each accession, two biological replications were used wherever possible for phenotyping and genotyping, although due to poor germination, four accessions (NMCA50034, PBC556, PBC1892, NMCA50064) did not have a biological replicate.

**Table 1.** *Capsicum* accessions included in this study

| Accession | Cultivar or other name | Species | Source |
| --- | --- | --- | --- |
| AVPP9905 | Susan's Joy | *Capsicum annuum* | WorldVeg |
| Criollo de Morelos 334 (CM334) | PBC 1867 | *Capsicum annuum* | NMSU |
| California Wonder | PBC 196 | *Capsicum annuum* | WorldVeg |
| PBC 1799 | Bird Pepper | *Capsicum annuum* | WorldVeg |
| VI012528 |  | *Capsicum baccatum* | WorldVeg |
| VI014924 | Aje | *Capsicum baccatum* | WorldVeg |
| PBC 80 |  | *Capsicum baccatum* | WorldVeg |
| PBC 81 | Jin's Delight | *Capsicum baccatum* | WorldVeg |
| NMCA90030 | PBC 1987 | *Capsicum cardenasii* | NMSU |
| NMCA90035 | PBC 1989 | *Capsicum cardenasii* | NMSU |
| PI 159236 | 30040 | *Capsicum chinense* | USDA-ARS |
| PI 152225 | Miscucho<br>Colorado | *Capsicum chinense* | USDA-ARS |
| VI012668 | PBC 306 | *Capsicum chinense* | WorldVeg |
| VI029446 |  | *Capsicum chinense* | WorldVeg |
| PBC 1793 | Scotch Bonnet<br>Pepper | *Capsicum chinense* | WorldVeg |
| VI013161 |  | *Capsicum eximium* | WorldVeg |
| VI012964 |  | *Capsicum eximium* | WorldVeg |
| NMCA90006 | PBC 1990 | *Capsicum eshbaughii* | NMSU |
| NMCA50030 | PBC 1991 | *Capsicum flexuosum* | NMSU |
| NMCA50034 | PBC 1992 | *Capsicum flexuosum* | NMSU |
| PBC 1820 | Bhut Jolokia | *Capsicum frutescens* × *chinense* | WorldVeg |
| PBC 556 | MC-003 | *Capsicum frutescens* | WorldVeg |
| NMCA50026 | PBC 1892 | *Capsicum galapagoensis* | NMSU |
| VI051011 |  | *Capsicum galapagoensis* | WorldVeg |
| NMCA50053 | PBC 1993 | *Capsicum minutifolium* | NMSU |
| NMCA90027 | PBC 1887 | *Capsicum praetermissum* | NMSU |
| VI029696 |  | *Capsicum praetermissum* | WorldVeg |
| VI029697 |  | *Capsicum praetermissium* | WorldVeg |
| NMCA50017 | PBC 1995 | *Capsicum rhomboideum* | NMSU |
| NMCA50064 | PBC 1996 | *Capsicum rhomboideum* | NMSU |
| VI051012 |  | *Capsicum tovarii* | WorldVeg |
| VI012478 |  | *Capsicum baccatum* | WorldVeg |
| VI059328 | PBC 142 | *Capsicum annuum* | WorldVeg |
| VI029657 |  | *Capsicum annuum* | WorldVeg |
| VI012900 |  | *Capsicum chacoense* | WorldVeg |
| PI 574547 | Chile que mira<br>p'arriba, PBC<br>1969 | *Capsicum annuum* var.<br>*glabriusculum* | USDA-ARS |
| PI 674459 | BG2816<br>selection 16-1,<br>PBC 1970 | *Capsicum annuum* var.<br>*glabriusculum* | USDA-ARS |
| VI012574 | PBC 814 | *Capsicum chacoense* | WorldVeg |

All experiments were conducted at the World Vegetable Center, Shanhua, Tainan, Taiwan (lat. 23.1°N; long. 120.3°E; elevation 12 m). Prior to sowing, all seed was treated with trisodium phosphate (TSP) and hydrochloric acid (HCl) following the methods of Kenyon et al. (2017), which has been observed to reduce germination rates. Seeds were sown into 72-cell plastic trays of sterilized peat moss. Trays were placed in a climate-controlled greenhouse for germination at 28 ± 3 °C with a 12-hour photoperiod and ≈95% relative humidity. At the 4-6 true leaf stage, the seedlings were transplanted into pots and moved to a greenhouse without climate control. Plants were irrigated twice daily and regularly fertilized with Nitrophoska (Incitec Pivot Fertilisers, Victoria, Australia) during the experimental period.

The accessions were morphologically characterized according to the Descriptors of *Capsicum* Manual (IPGRI et al., 1995) for the following characteristics: mature leaf length, mature leaf width at widest point, leaf color, density (if present) of leaf pubescence, leaf shape, lamina margin, stem color, stem shape, density (if present) of stem pubescence, nodal anthocyanin color, node length, anther color, anther length, filament length, corolla color, corolla spot color, corolla shape, corolla length, stigma exsertion, flower position, tillering, leaf density, fruit length, fruit width, fruit pedicel length, neck at base of fruit. Quantitative traits were the mean of 10 values measured across replicates. Qualitative traits were scored according to the IPGRI Descriptors of *Capsicum* manual based on observations of both plant replicates. Accessions with incomplete data were excluded from analysis of phenotypic data. To identify trends in traits between species, the quantitative traits were used for principal component analysis (PCA) using the R packages, ‘factoextra’ (Kassambara and Mundt, 2020) and ‘ggfortify’ (Horikoshi et al., 2020) for PCA analysis with scaling. The scores of qualitative traits were analyzed using an unweighted pair group method with arithmetic mean (UPGMA) hierarchical cluster analysis. Bootstrap resampling was applied to clustering with 1,000 iterations.

Reciprocal hybridizations were attempted among all combinations of accessions throughout the experimental period. Ability to hybridize in reciprocal was used to confirm previous reports of relatedness and ability to hybridize species across clades and complexes. The fruits of successful hybridizations were collected upon ripening. Within three days of harvest, the seeds were extracted from the fruits and dried for at least 1week. Five seeds each of 112 crosses of interest were sown into 70-cell plastic trays containing sterilized peat moss. The trays were placed in a greenhouse without climate control and irrigated twice daily and observed daily for 12 weeks to assess germination. A chord diagram was produced in R using the package ‘circlize’ (Gu et al., 2014) to visualize successful crosses for which seed was obtained. A heat map was produced in R using the package ‘ggplot2’ (Wickham et al., 2020) to visualise the percentage of seeds germinated after 12 weeks.

For genotyping, DNA was isolated from young, actively growing leaves from plants of each accession using the modified cetrimonium bromide (CTAB) extraction method (Meyer and Buchta, 2020). Using 27 Simple Sequence Repeat (SSR) markers, DNA was amplified by PCR, for which each well of a 96-well microtiter plate contained 2 μl of template DNA, 0.4 μl of primer (0.2 μl each forward and reverse), 0.1 μl of AmpliTaq Gold DNA polymerase, 0.4 μl of deoxyribonucleotides, 1.5 μl of 10× PCR Buffer II Gold buffer (Thermo Fisher Scientific, Waltham, MA, USA), and sterile water to a final volume of 15 μl. The reactions were carried out in a thermal cycler (Single Block Alpha Unit, DNA Engine^®^, Bio-Rad Laboratories, Berkeley, CA, USA) with an annealing temperature of 55 °C. The electrophoresis of amplified products was performed on 6% acrylamide gels at 160 Volts for 30 minutes (Thermo Electron Electrophoresis EC250-90, Thermo Fisher Scientific). The results were visualized under UV light using UVITEC Imaging Systems (Cleaver Scientific, Warwickshire, UK) following staining with ethidium bromide. Electrophoresis was repeated whenever the clarity of the bands or their exact size was uncertain.

Gels were scored for each primer pair using a binary method: each accession was scored for presence (1) or absence (0) of amplicons of each size. The data were processed in R using the packages, ‘proxy’ (Shipunov et al., 2020) and ‘shipunov’ (Altschul et al., 1990) to produce a dendrogram with bootstrapping for the assessment of the relatedness between the individual accessions. A distance matrix was produced using the Dice index, and an unweighted pair group method with arithmetic mean (UPGMA) hierarchical cluster analysis was carried out. Bootstrap resampling was applied to clustering with 1,000 iterations.

Further molecular analysis to clarify the identification of some accessions included the study of the *waxy* gene region of six accessions, VI051012 (*C. tovarii*); VI051011 (*C. galapagensis*, potentially *C. annuum*); VI012574 (*C. chacoense*, potentially *C. annuum*); PBC1892 (*C. galapagensis*); VI013161 (*C. eximium*); and PBC 556 (*C. frutescens*), using the primer pair, 860F and 2R (Carrizo García et al., 2016). The chosen accessions were those expected to need clarification due to possible misidentification, based on molecular and morphological data. The *waxy* region was amplified by PCR as before, with an annealing temperature of 60 °C. The quality of the products were evaluated by running on a 2% agarose gel with EtB‘out’ (Yeastern Biotech Co. Ltd., Taipei, Taiwan) at 100 Volts for 50 minutes, then visualized using a Microtek Bio- 1000F gel imager (Microtek International Inc., Hsinchu, Taiwan). The PCR products were sequenced by Genomics Biotechnology Co., Ltd. (New Taipei City, Taiwan) by the Sanger sequencing method. Low-quality nucleotides were manually removed throughout the resulting sequence, including approximately the first and last 60 nucleotides. The sequences were aligned using NCBI nucleotide BLAST (Hall, 1999) and a consensus sequence constructed using the CAP contig assembly program from BioEdit (Katoh and Standley, 2013). The sequences were aligned using multiple sequence alignment tool, Clustal MAFFT (Katoh and Standley, 2013), to produce a dendrogram.

## Results

To clarify the phylogeny of the wild and domesticated *Capsicum* species in the sample, UPGMA clustering was carried out using the SSR molecular markers. The *C. baccatum* and *C. chinense* group accessions were distinct from the *C. annuum* group, with 76% bootstrap support (Fig. 1). The *C. baccatum* accessions made up a significant group, being closely clustered with the *C. praetermissium* accessions, as expected. This grouping was adjacent to a large group comprised of closely clustered *C. chinense* accessions with the *C. galapagensis* accession PBC1892, as well as *C. eshbaughii*, *C. eximium* and *C. frutescens*, separated from the *C. baccatum* group with a relatively low confidence interval. Within this grouping, *C. chinense* accession, PBC1820, was distinct from its counterparts, supported by 92% bootstrap support. Furthermore, the *C. chinense* species accessions were relatively separate from the accessions at the periphery of this grouping, the *C. galapagensis* accession PCB1892, the *C. frutescens* accession PBC556, and the *C. eximium* accession VI013161. The grouping of the *C. eximium* accession VI013161 with *C. frutescens* was similar in clustering from the *waxy* gene sequence (Fig. 2). These accessions were thus more similar to each other than they were similar to the *C. galapagensis* accession PBC1892, and this was a distinct grouping from other sequenced accessions.

**Figure 1.**
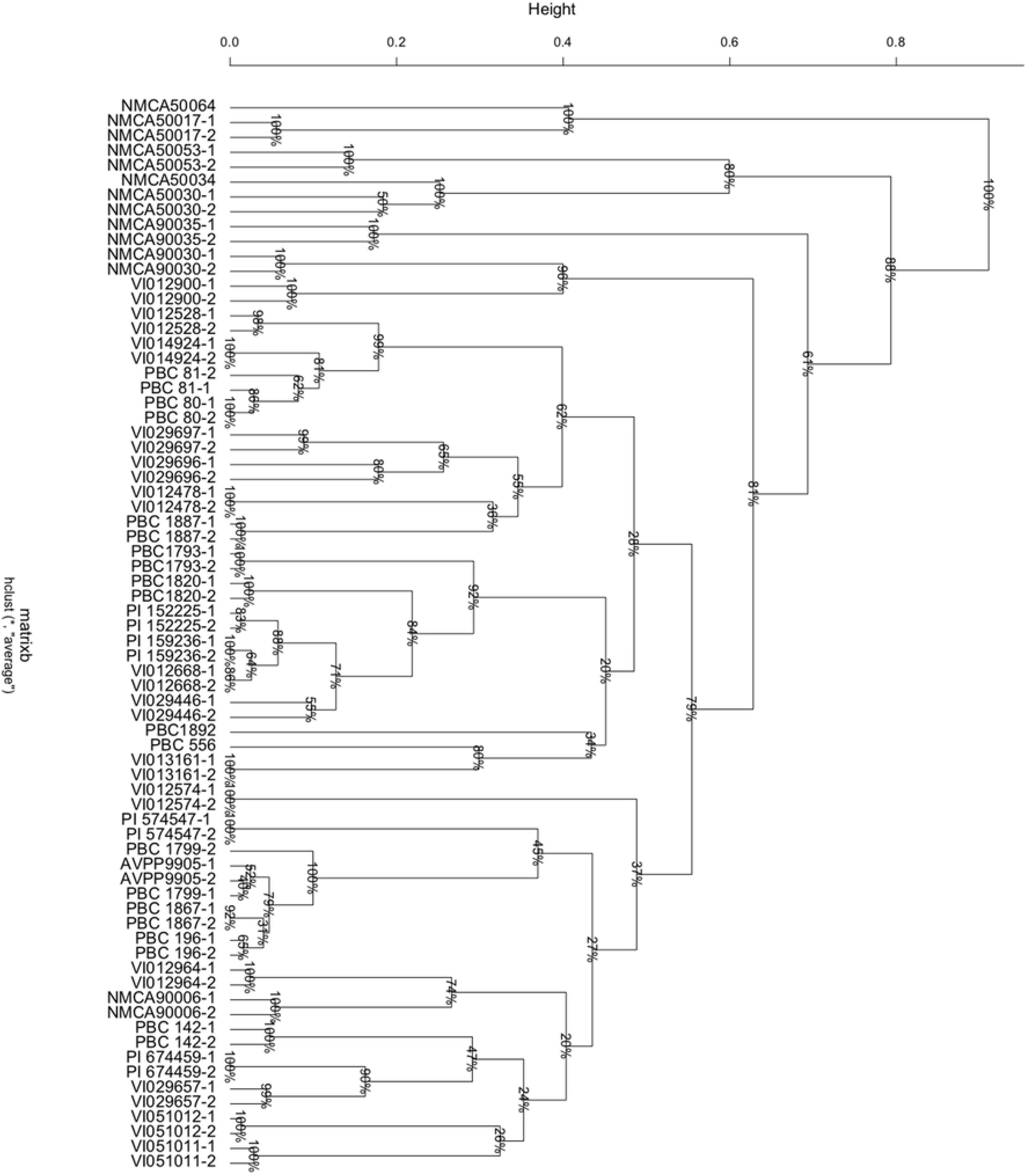
Unweighted pair group method (UPGMA) clustering of *Capsicum* species according to single sequence repeat (SSR) markers. ‘Height’ represents dissimilarity, derived from ‘dice method’. Bootstrap resampling applied to clusters, represented as percent confidence interval. Numbers following the hyphen indicate replicates.

**Figure 2.**
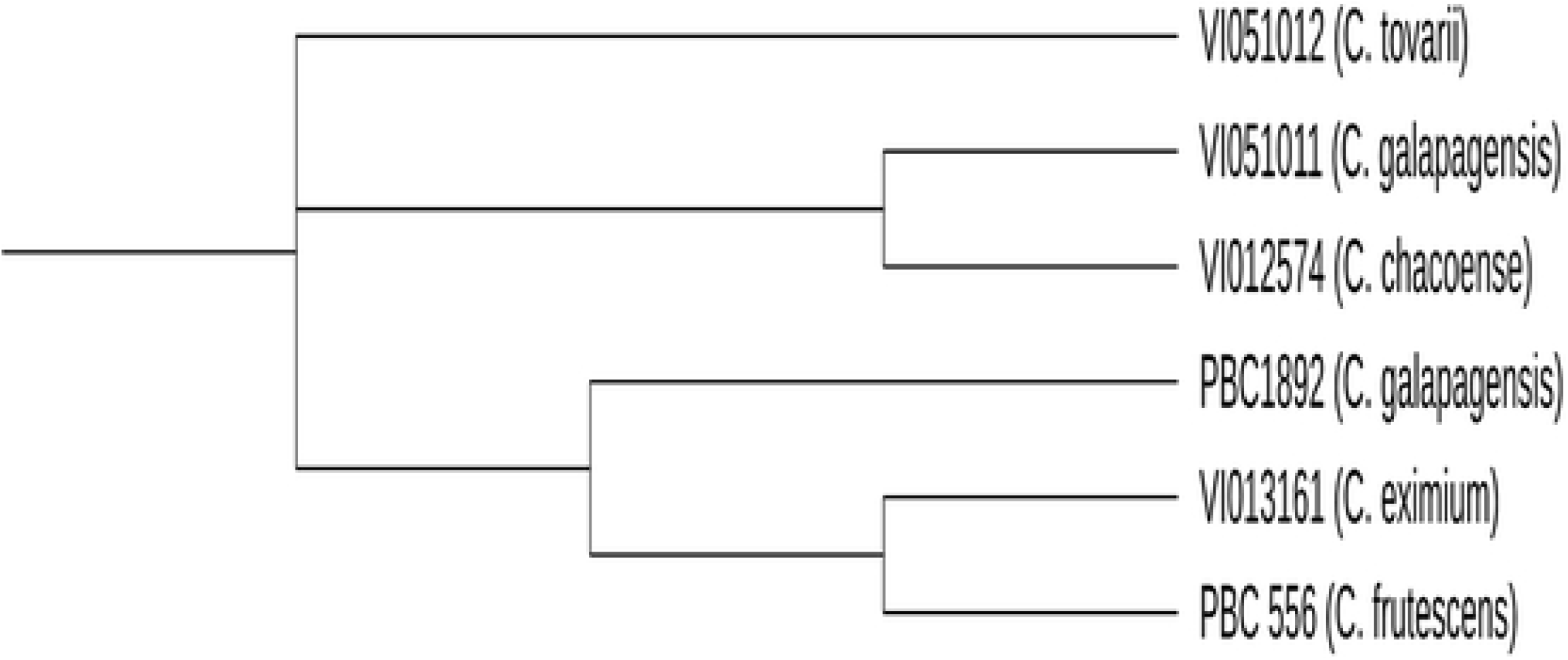
Clustering of selected *Capsicum* species according to their *waxy* gene sequences.

To better understand the crossing relationship between species, reciprocal hybridizations were performed between each combination of accessions (Fig. 3). Members of *C. baccatum* and *C. praetermissium* hybridized as either the female or male parent with at least one accession of each other species with the exception of *C. rhomboideum* (Fig. 3). However, of the sample of seeds selected for sowing, only the cross between VI014924 and PBC1969 germinated (Fig. 4). Hybridizations were not achieved between *C. galapagoensis* as either parent with accessions of *C. tovarii*, *C. flexuosum*, *C. minutifolium*, *C. cardenasii*, *C. eshbaughii*, and *C. rhomboideum* species. *Capsicum eshbaughii* hybridized more readily as the female parent, but failed to hybridize in either direction with accessions of *C. eximium*, *C. frutescens*, *C. galapagoensis*, *C. tovarii*, *C. flexuosum*, and *C. rhomboideum*. The majority of *C. frutescens* hybrids were achieved with *C. annuum* accessions, but successful hybridizations were found across a broad species range. Of the sample of seeds sown, 80% of the PBC556 × PBC1970 cross seeds germinated (Fig. 4).

**Figure 3.**
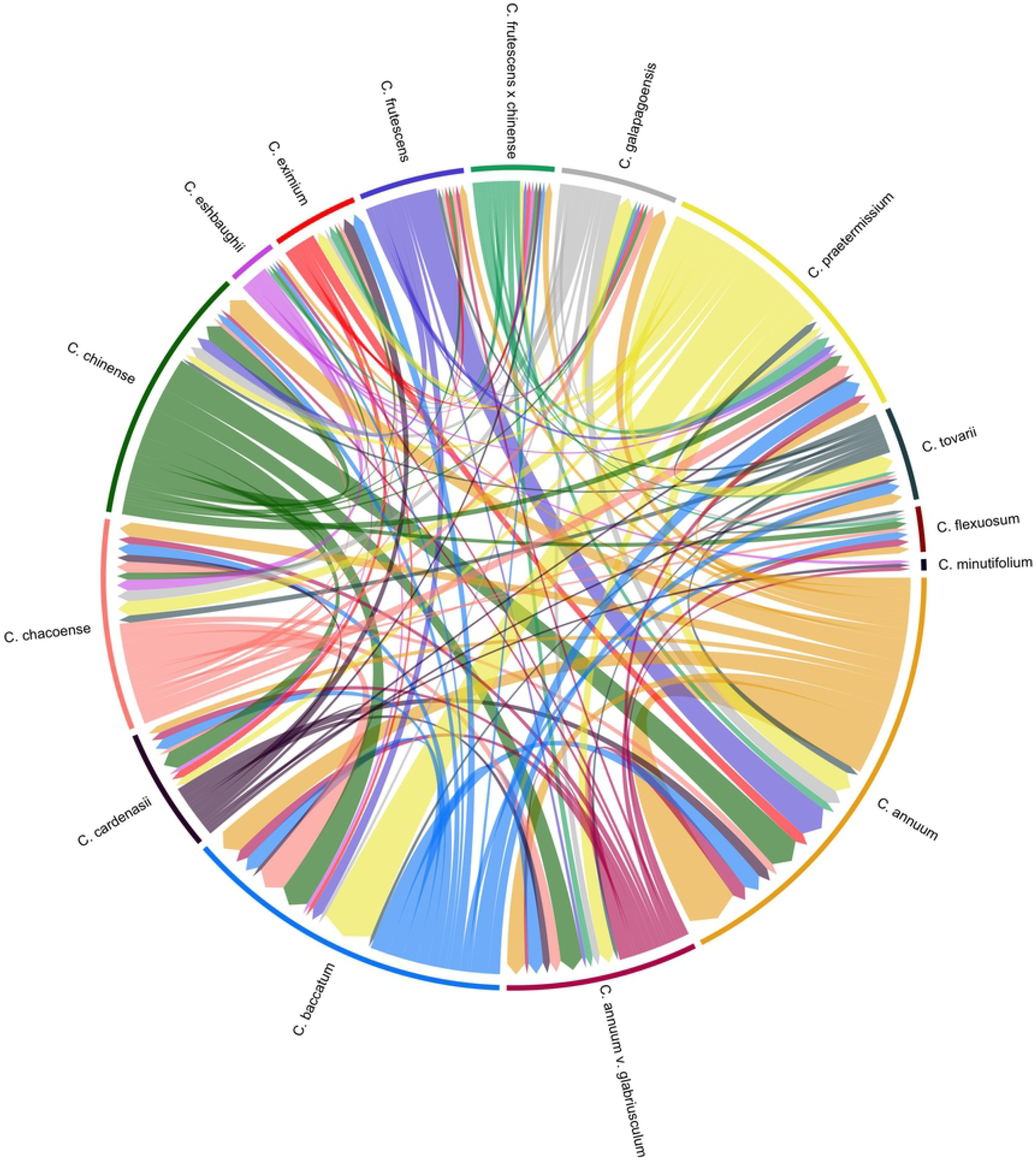
Reciprocal hybridizations achieved between accessions of *Capsicum* species. Direction of arrow represents successful hybridizations in the male-female direction from which fruit was harvested.

**Figure 4.**
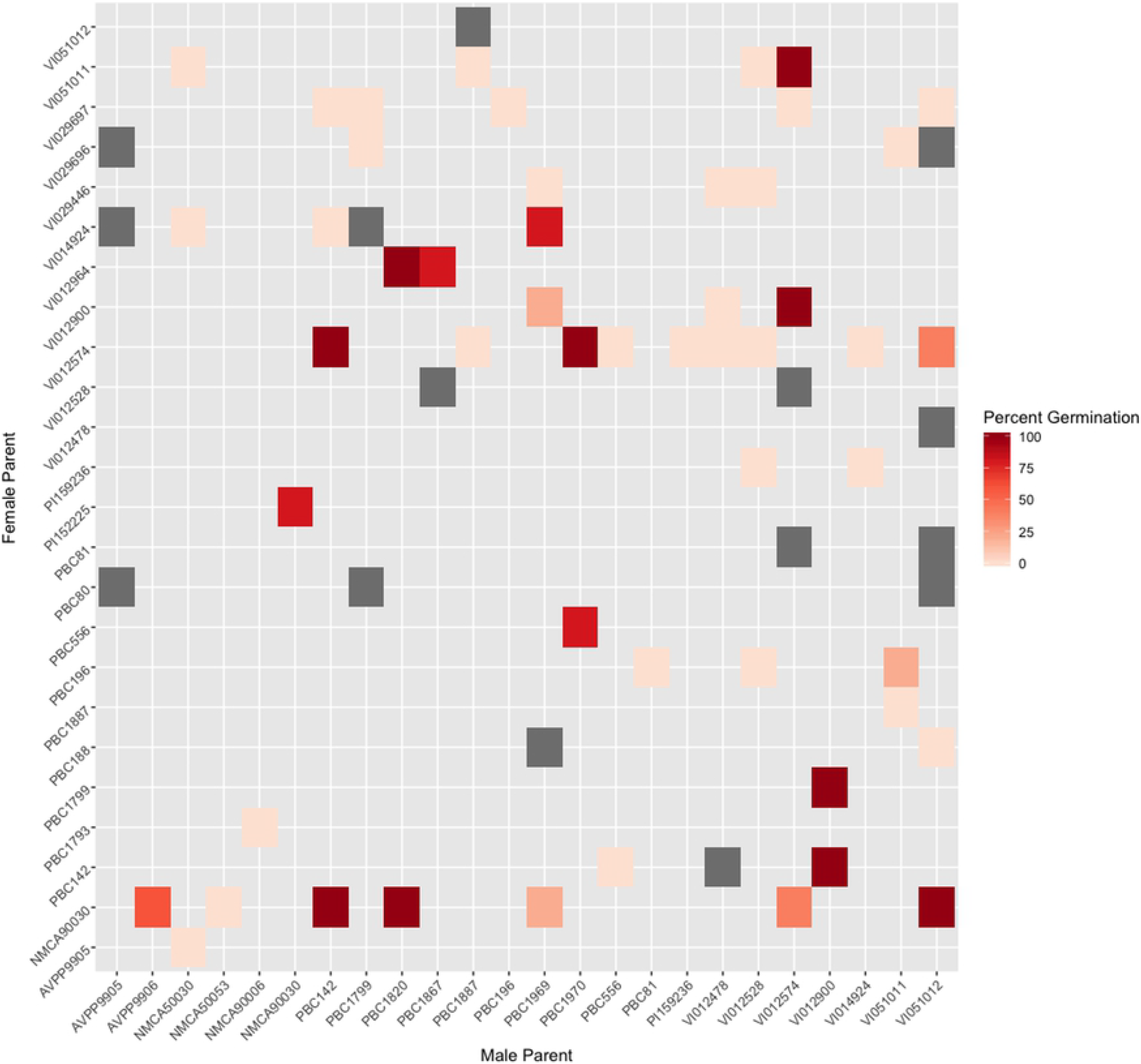
Percent germination of selected hybrid seeds 12 weeks after sowing. Grey indicates unviable seeds.

The *C. annuum* group, which was adjacent to *C. baccatum*, consisted of the closely clustered *C. annuum* species accessions: PBC1799, AVPP9905, PBC1899, PBC1867, and PBC196, along with *C. annuum* var. *glabriusculum* PI574547, and *C. chacoense* VI012574 (Fig. 1). Neighboring this group was a cluster comprised of the *C. eximium* accession VI012964, the *C. eshbaughii* accession NMCA90006, the *C. annuum* accessions PBC142 and VI029657, the *C. annuum* var. *glabriusculum* accession PI674459, the *C. tovarii* accession VI051012, and the *C. galapagoensis* accession VI051011. Based on clustering of the *waxy* gene sequence, we found *C*. *tovarii* to be less similar to *C. galapagoensis* and *C. chacoense* than they were to one another (Fig. 2). Furthermore, the *C. galapagoensis* accession VI051011 and the *C. chacoense* accession VI02574 were similar in the *waxy* gene region, supporting their grouping together adjacent to the *C*. *annuum* group.

Accessions of *C. annuum* hybridized in both directions with one or more accessions of all species except *C. rhomboideum* and *C. minutifolium* (Fig. 3). Thirteen of the hybrids sown germinated well (Fig. 4). *Capsicum annuum* var. *glabriusculum* hybridized in either direction with at least one accession of every species except *C. rhomboideum*, and 7 out of 10 of those sown germinated (Fig. 4). Accessions of *C*. *chacoense* also hybridized broadly, but not with *C. frutescens* × *chinense, C. minutifolium* or *C. rhomboideum*, and seven of the 25 hybrids sown germinated. More than one cross was achieved between *C. tovarii* and an accession of every species except *C. eshbaughii*, *C. eximium*, *C. galapagoensis*, *C. minutifolium* and *C. rhomboideum*. Of these crosses, VI012574 × VI051012 and NMCA90030 × VI051012 germinated with 40% and 100% efficiency, respectively.

With 80% confidence interval, *C. chacoense* accession VI012900 and *C. cardenasii* accession NMCA90030 were clustered separately from the *C. baccatum* and *C. annuum* groups (Fig. 1). NMCA90035 clustered distinctly from its *C. cardenasii* counterpart, with bootstrap support of 57%. The *C. flexuosum* accessions NMCA50034 and NMCA50030 clustered closely together, with high bootstrap support (99%); adjacent was the *C. minutifolium* accession NMCA50053, and in a separate cluster, the *C. rhomboideium* accession NMCA50064, which was the most distinct grouping, separated from its neighbors with 100% confidence.

*Capsicum cardenasii* species hybridized with every species except *C. frutescens, C. flexuosum, C. galapagensis* and *C. rhomboideum* (Fig. 3), and the five of the nine hybrids sown germinated well (Fig. 4). No hybrids were achieved with *C. flexuosum* or *C. minutifolium* as female parents, however *C. flexuosum* hybridized as the male parent with *C. chacoense*, *C. annuum* var. *glabriusculum*, *C. baccatum*, *C. tovarii*, *C. annuum* and *C. frutescens*, while *C. minutifolium* hybridized with *C. cardenasii*, *C. eshbaughii*, and *C. annuum* var. *glabriusculum*, and of the crosses sown, only VI012574 × PBC124 germinated (Fig. 4). No successful hybrids were achieved with *C. rhomboideum* in either direction (Fig. 3).

We applied principal component analysis to the quantitative phenotypic data collected to understand phenotype across *Capsicum* species (Fig. 5). The first two components account for 59.9% of the total variation. The *C. baccatum* accessions made up a group along with PBC196 and VI01668, due to their correlated fruit and flower characteristics (pedicel length, fruit width, fruit length, anther length, filament length, corolla length) (Fig. 5). *Capsicum annuum* accessions made up a less distinct group, along with the wild progenitor *C. annuum glabriusculum*, and the other domesticated species *C. chinense*, *C. frutescens*, *C. frutescens* × *chinense* along with *C. eximium* (Fig. 5).

**Figure 5.**
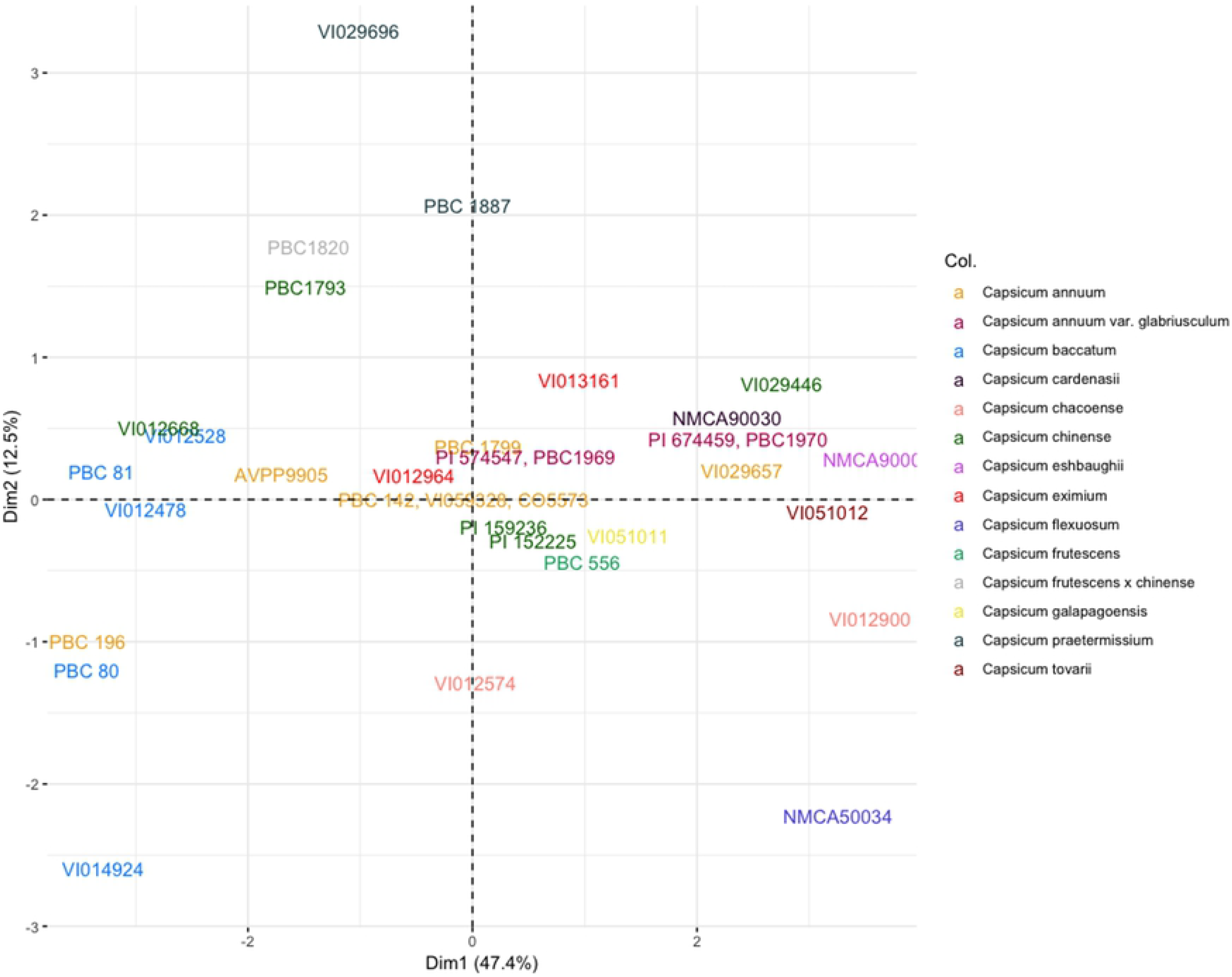
First two principal components of accessions in the wild and domesticated *Capsicum* species based on the quantitative phenotypic data.

The UGMA clustering of the accessions’ qualitative phenotypic data more closely mirrored the genetic relatedness based on SSR molecular markers, especially for the *C. baccatum* and *C*. *praetermissum* accessions (Fig. 6), which formed two groupings (VI012528, VI029696, VI029697; and PBC81, VI014924, PBC81). However, we found *C. annuum* did not form a unique clade, highlighting the phenotypic diversity of this domesticated species (Fig. 6).

**Figure 6.**
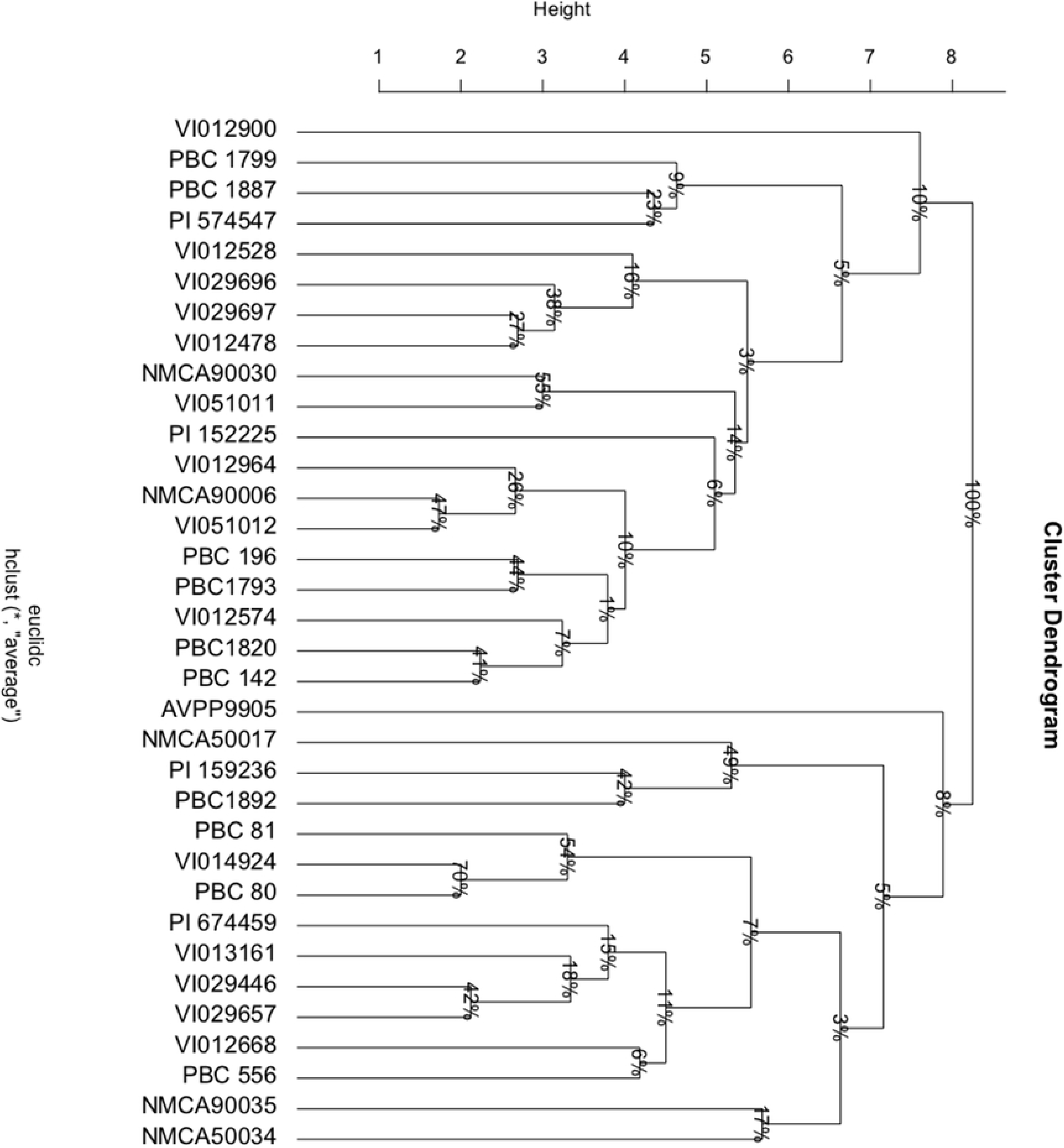
Unweighted pair group method (UPGMA) clustering of wild and domesticated *Capsicum* species based on the qualitative phenotypic data, scored according to IPGRI *Descriptors of Capsicum* scoring method. Bootstrap resampling applied to clusters, represented as percent confidence interval.

## Discussion

Understanding the relatedness between accessions of *Capsicum* species, and the extent to which they hybridize is key in identifying candidates for the introgression of traits of interest into commercial varieties. Our results mirror the widely accepted species phylogeny, with clustering based on SSR markers centered around *C. annuum*, *C. baccatum* and *C. chinense* complexes (Bosland and Votava, 2012; Pickersgill, 1971; Emboden Jr., 1962). Similarly, our results of phenotyping the accessions also support previously described genetic complexes (Carrizo García et al., 2016). Generally, accessions from the domesticated species (*C*. *annuum*, *C*. *chinense*, and *C*. *frutescens*) were nearer the origin of the score plot, with the exception of members of the domesticated *C*. *baccatum*, which clustered further away from the other domesticated species (Fig. 5). Conversely, members of the wild species were further away, indicating greater diversity (Fig. 5). Although we measured different phenotypic traits, our findings contradict those of Luna-Ruiz et al. (2018), who found greater levels of diversity among domesticated species for capsaicinoids. Interestingly, based on hybridization success rates, there was a weak relationship between relatedness and crossibility, which is in contrast with previous understanding that compatibility between complexes is low (van Zonneveld et al., 2015). This suggests potential for crop improvement with wild relatives of domesticated species with genetic bridge strategies.

When phylogeny, interspecific compatibility and phenotype are considered in concert, the identity of a number of accessions included in this study may be questioned. The issue of misidentification of *Capsicum* species has been raised previously, with several genebanks incorrectly reporting accessions *C. frutescens* as *C. galapagoense* (P.W. Bosland, pers. comm.). Thorough characterization is important in supporting conservation of genetic material and identifying gaps in genebank collections (FAO, 2010). Only 12% of national vegetable germplasm collections have been characterized biochemically, while 65% have been characterized morphologically (FAO, 2010). Thorough characterization is therefore key in understanding the reproductive relationships between *Capsicum* species.

Clustering based on SSR genotyping revealed a close relationship between *C. baccatum* accessions (Fig. 1), as expected for this well-accepted domesticated species. *Capsicum praetermissium* accessions are also grouped within this complex, which supports previous findings (Albrecht et al., 2012) and suggestions that *C. praetermissium* in fact comprises a subgroup of *C. baccatum* (Albrecht et al., 2012). Both PCA and UPGMA analysis of *C. baccatum* and *C. praetermissium* morphologies also show their close clustering, further demonstrating their relationship. *Capsicum praetermissium* is thought to have diverged prior to domestication of *C*. *baccatum*, and has not been utilized in breeding domestic *C. baccatum* accessions (Albrecht et al., 2011). We found *C. praetermissium* readily hybridized with *C. baccatum* (Fig. 3) in line with the findings of Emboden Jr. (1961), and thus offers potential as a genetic resource.

*Capsicum chinense* species accessions comprised a significant cluster, which included *C. chinense*, *C. frutescens*, *C. eshbaughii*, and *C. galapagoensis* (Fig. 1). The grouping of *C. chinense* adjacent to *C. baccatum* was in line with a recent study that also used SSR molecular markers to characterize *Capsicum* species (Guzmán et al., 2020). However, this is contrary to findings of Pickersgill et al. (1991) and Ince et al. (2010), who grouped *C. chinense* within the *C. annuum* complex. Furthermore, in this study, a total of 20 crosses were achieved between *C. annuum* (including the wild progenitor *C. annuum* var. *glabriusculum*), and *C. chinense*, 13 of which had a *C. chinense* female parent (Fig. 3). Seeds from two of these crosses were sown (VI029446 × PBC1969 and PI152225 × NMCA90030) and germinated well (Fig. 4). This contrasts to previous work that reports a barrier to reproduction between *C. annuum* and *C. chinense* (Campos et al., 2004). However, Costa et al., (2009) found that crosses between *C. chinense* and *C. annuum* accession were possible. This highlights the variability in compatibility, and its dependence on accession selection.

The grouping of *C. frutescens* in the *C. chinense* complex was in line with previous findings of the close relationship of these species (Fig. 1) (Shiragaki et al., 2020). A number of researchers argued their identities as sister species within the annuum clade (Albrecht et al., 2012; Tam et al., 2009; Walsh and Hoot, 2001). Furthermore, we found *C. frutescens* hybridized readily with both members of the *C. baccatum* and *C. annuum* clades, as well as with *C. chinense* (Fig. 3). Of the three *C. frutescens* hybrids selected for sowing, 80% of the PBC556 × PBC1970 hybrid seeds germinated (Fig. 4). The relationship of *C. eshbaughii* to this clade, and its pairing with *C. eximium* was consistent with its previous placement in the ‘Purple Corolla clade’ (Carrizo García et al., 2016). *Capsicum chinense* formed hybrids with other members of this grouping (Fig. 3), and 100% of seeds from the *C. chinense* and *C. eximium* cross germinated (Fig. 4).

Interestingly, *C. eximium* and *C. cardenasii* appeared distantly related in our analysis (Fig. 1), which contradicts previous reports of these species being within the *C. pubescens* complex (Ibiza et al., 2012; McLeod et al., 1983). Furthermore, their phenotypes correlated closely with *C. annuum* accessions (Fig. 5). This raises the question of the validity of the identification of accessions VI013161 and VI012964 as *C. eximium*.

The *C. annuum* accessions comprise a major grouping adjacent to the *C. baccatum* group (Fig. 1). A sample of *C. annuum* accessions (PBC1799, PBC196, PBC1867 and AVPP9905) formed a tightly clustered group, indicating genetic similarity. They also display highly correlated phenotypes, forming a cluster along with accessions from other domesticated species (Fig. 5). The *C. annuum* accessions PBC142 and VI029657 were in an adjacent group (Fig. 1), therefore may be considered part of the wider *C. annuum* complex, along with *C. chacoense* accession VI012574, *C. galapagensis*, VI051011, and *C. tovarii* accession VI051012. The presence of *C. chacoense* (VI012574) in this group, distant from the second *C. chacoense* accession (VI012900) included in this study, highlights its possible misidentification. Sequencing clustered VI012574 with *C. galapagensis* accession VI051011, which may be considered a member of the *C. annuum* complex (Fig. 2). Principal component analysis (Fig. 5) revealed VI012574 was grouped with *C. annuum* accessions, away from its counterpart, while UPGMA analysis further highlights this disparity. Direct observation of the phenotypes emphasises the similarity between the morphology of VI012574 and typical *C. annuum* features. This includes upright growth, elongated fruits, and relatively large flowers with blue anthers.

The *C. galapagoensis* accession, PBC1892 was grouped with the wider *C. baccatum* cluster (Fig. 1), conflicting previous findings that *C. galapagoensis* is derived from a *C. annuum* progenitor population (Choong, 1998). No successful hybridizations were achieved between PBC1892 and PBC556 (*C. frutescens*), which clustering suggested were closely related. The second *C. galapagoensis* accession included in the study, VI051011, was grouped within the *C. annuum* complex, and displayed a distinctly different phenotype to that of PBC1892. PBC1892 had a compact growth habit, very small fruits, flowers and leaves, and densely pubescent stems and leaves, typical of *C. galapagoensis* descriptions. Conversely, VI051011had a morphology similar to that of *C. annuum*, reflected in its close proximity to the PCA origin, along with *C. annuum* accessions. Eight hybridizations were achieved between VI051011 and *C. annuum* accessions, and of the selected hybrid seeds sown, 20% germinated. The close clustering of VI051011 with the *C. annuum* complex, their similar morphology and their ability to hybridize suggests likely misidentification of this accession.

The grouping of VI051011 (*C. galapagoensis*) in the *C. annuum* complex (Fig. 1) supports previous findings that *C. galapagoensis* is derived from a *C. annuum* progenitor population (Choong, 1998). Eight hybridizations were achieved between VI051011 and *C. annuum* accessions, and of the selected hybrid seeds sown, 20% germinated. The second *C. galapagoensis* accession in this study, PBC1892, was grouped with the wider *C. baccatum* cluster (Fig. 1). No successful hybridizations were achieved between PBC1892 and PBC556 (*C. frutescens*), which clustering suggested were closely related. The phenotypes of these accessions were distinctly different; PBC1892 having a compact growth habit, very small fruits, flowers and leaves, and densely pubescent stems and leaves, which is typical of *C. galapagoense* descriptions. VI051011 had morphology similar to that of *C. annuum* (Fig. 5) and its grouping within the *C. annuum* complex along with its ability to hybridize with *C. annuum* suggested likely misidentification.

There were five further clusters consisting of *C. chacoense, C. cardenasii, C. flexuosum, C. minutifolium*, and *C. rhomboideum* respectively, which had increasingly distant relation to the three major species complexes (Fig. 1). Although *C. chacoense* has been previously grouped within the *C. baccatum* clade (Carrizo García et al., 2016; McLeod et al., 1983), this wild species has an apparently distant relationship with *C. baccatum. Capsicum cardenasii* was similarly distantly related to other clades. Other studies (Ibiza et al., 2012; Ince et al., 2010; McLeod et al., 1983) also found *C. chacoense* and *C. cardenasii* not to be closely related to any major clade. Furthermore, VI012900 hybridized readily with members of both *C. annuum* and *C. baccatum* clades (Fig. 3). Furthermore, both *C. chacoense* and *C. cardenasii* accessions (with the exception of PI159236 and PI15225) lay on the periphery of the PCA plot, clustering with neither *C. baccatum* or *C. annum* groups. . This suggests *C. cardenasii* and *C. chacoense* accessions are not members of either *C. baccatum* or *C. annuum* clades.

*Capsicum flexuosum, C. minutifolium* and *C. rhomboideum* were distantly related to the major clades, consistent with the body of literature (Carrizo García et al., 2016; Guzmán et al., 2020; Choong, 1998). The *C. flexuosum* accession (NMCA50034) also had a distant relationship with more domesticated species (Fig. 5). A small number of hybridizations were achieved between *C. flexuosum* and *C. minutifolium* with members of both *C. annuum* and *C. baccatum* clades. However, no hybridizations were achieved between *C. rhomboideum* and any other accession.

The results reported here highlight the extent of phenotypic diversity in *Capsicum* species, the complexity of *Capsicum* phylogeny, and the similarly complex reproductive relationships between *Capsicum* species. The evidence suggesting the incorrect identification of VI013161, VI012964, VI012574, and VI051011 may highlight a broader issue of misidentification of *Capsicum* in genebanks. Thorough characterization of *Capsicum* genetic material taking a multifaceted approach is therefore important for the development of future breeding programs. Furthermore, the generation of diverse hybrids among accessions of all species included in this study (with the exception of *C. rhomboideum*) demonstrates the possibility for introgression of a diverse range of traits of interest directly or through the design of bridge crossing strategies. Wild relatives of domesticated *Capsicum* species therefore represent significant potential for future breeding programs, and should not be discounted on the basis of their assumed relatedness to domesticated species.

## Acknowledgments

We thank Dr. Paul Bosland of Chile Pepper Institute, New Mexico State University, USA for providing *Capsicum* accessions.

## Copyright

© 2021 Parry et al. This is an open access article distributed under the terms of the <u>Creative Commons Attribution License</u>, which permits unrestricted use, distribution, and reproduction in any medium, provided the original author and source are credited.

## Data Availability

The data used in this study are available in the World Vegetable Center repository, HARVEST, doi:10.22001/wvc.73914, https://worldveg.tind.io/record/73914

## Literature Cited

Aguilar Meléndez A, Morrell PL, Roose ML, Kim SC. Genetic diversity and structure in semiwild and domesticated chiles (*Capsicum annuum*; Solanaceae) from Mexico. Amer J Bot. 2009;96(6):1190–1202.

Albrecht E, Zhang D, Mays AD, Saftner RA, Stommel JR. Genetic diversity in *Capsicum baccatum* is significantly influenced by its ecogeographical distribution. BMC Genet 2012;13 https://doi.org/10.1186/1471-2156-13-68.

Albrecht E, Zhang D, Saftner RS, Stommel JR. Genetic diversity and population structure of *Capsicum baccatum* genetic resources. Gene Resources Crop Evol. 2011;59(4):517–38. https://doi.org/10.1007/s10722-011-9700-y.

Altschul SF, Gish W, Miller W, Myers EW, Lipman DJ. Basic local alignment search tool. J Molec Biol. 2009;215(3)403–10. https://doi.org/10.1016/S0022-2836(05)80360

Barchenger DW, Naresh P, Kumar S. Genetic resources of *Capsicum*. In: Ramchiary N, Kole C, editors. The Capsicum Genome. New York: Springer Nature; 2019; p. 9–23.

Barchenger DW, Bosland PW. Wild chile pepper (*Capsicum* L.) of North America. In: Greene S, Williams K, Khoury C, Kantar M, Marek L, editors. North American Crop Wild Relatives, New York: Cham: Springer International Publishing; 2019; p. 225–42.

Bosland PW. Sources of curly top virus resistance in *Capsicum*. HortScience 2000;35(7):1321–2.

Bosland PW, Votava EJ. Peppers: Vegetable and spice capsicums. 2nd ed. 2012, Wallingford. CABI

Campos KP, Pereira TNS, Costa FR, Sudré CP, Monteiro CES, Rodrigues R. Interspecific Hybridization among cultivated germplasm in *Capsicum*. 2004 Naples, Florida, 17th International Pepper Conference. p.20.

Carrizo García C, Barfuss MHJ, Sehr EM, Barboza GE, Samuel R, Moscone EA, et al. Phylogenetic relationships, diversification and expansion of chili peppers (*Capsicum*, Solanaceae). Ann Bot, 2016;118(1):35–51. https://doi.org/10.1093/aob/mcw079.

Cheng J, Qin C, Tang X, Zhou H, Hu Y, Zhao Z, et al. Development of a SNP array and its application to genetic mapping and diversity assessment in pepper (*Capsicum* spp.). Sci Rep, 2016;6:1–11 https://doi.org/10.1038/srep3329.

Choong CY. DNA Polymorphisms in the study of relationships and evolution in *Capsicum* [dissertation]. Reading, UK: Univ. Reading; 1998.

Costa LV, Lopes R, Lopes MTG, de Figueiredo AF, Barros WS, Alves RRM. 2009. Cross compatibility of domesticated hot pepper and cultivated sweet pepper. Crop Breed Appl Biotechnol. 2014;9:37–44.

Eggink PM, Tikunov Y, Maliepaard C, Haanstra JP, de Rooij H, Vogelaar A, et al. Capturing flavors from *Capsicum baccatum* by introgression in sweet pepper. Theor Appl Genet. 2014;127(2):373–390. https://doi.org/10.1007/s00122-013-2225-3.

Emboden Jr. WA. A preliminary study of the crossing relationships of *Capsicum baccatum*. Butler Univ Bot Stud. 1962;14:108–114. https://doi.org/10.1080/00231940.1962.11757638.

FAO. The second report on the state of the world’s plant genetic resources for food and agriculture. FAO, Rome; 2010 http://www.fao.org/3/i1500e/i1500e00.htm

Gramazio P, Prohens J, Plazas M, Mangino G, Herraiz FJ, Vilanova S. Development and genetic characterization of advanced backcross materials and an introgression line population of *Solanum incanum* in a *S. melongena* background. Frontiers Plant Sci. 2017;8:1477. doi:10.3389/fpls.2017.01477

Gu Z, Gu L, Eils R, Schlesner M, Brors B. *circlize* Implements and enhances circular visualisation in R. Bioinformatics. 2014;30(19):2811–2812 https://doi.org/10.1093/bioinformatics/btu393.

Guzmán FA, Moore S, de Vicente MC, Jahn MM. Microsatellites to enhance characterization, conservation and breeding value of *Capsicum* germplasm. Genet Resour Crop Evol. 2020;67(3):569–585. https://doi.org/10.1007/s10722-019-00801-w.

Hall TA. Bioedit: a used-friendly biological sequence alignment editor and analysis program for Windows 95/98/NT. Nucleic Acids Symp Ser. 1999;4:195–98 https://doi.org/10.14601/PHYTOPATHOL_MEDITERR-14998U1.29.

Hirsch CN, Hirsch CD, Felcher K, Coombs J, Zarka D, van Deynze A, et al. Retrospective view of North American potato (*Solanum tuberosum* L.) breeding in the 20th and 21st centuries. G3-Genes, Genom, Genet. 2013;3(6):1003–1013. https://doi.org/10.1534/g3.113.005595.

Horikoshi M., Tang Y, Dickey A, Genié M, Thompson R, Seltzer L, et al. ggfortify: Data visualization tools for statistical analysis results. R package version 0.4.11. 2020

Ibiza VP, Blanca J, Cañizares J, Nuez F. Taxonomy and genetic diversity of domesticated *Capsicum* species in the Andean region. Genet Resour Crop Evol., 2012;59(6):1077–1088. https://doi.org/10.1007/s10722-011-9744-z.

Ince AG, Karaca M, Onus AN. Genetic relationships within and between *Capsicum* species. Biochem Genet. 2010;48(1-2):83–95. https://doi.org/10.1007/s10528-009-9297-4.

IPGRI, AVRDC and CATIE. Descriptors for Capsicum (Capsicum spp.). International Plant Genetic Resources Institute, Rome, Italy; the Asian Vegetable Research and Development Center, Taipei, Taiwan, and the Centro Agronómico Enseñanza, Turrialba, Costa Rica; 1995.

Jimenez RC. Utilizing Wild *Capsicum annuum* germplasm for breeding resistance to beet curly top virus (genus: *Curtovirus*, family: *Geminiviridae*) in Cultivated Pepper (*Capsicum annuum* L.) [PhD Thesis]. University of California, Davis; 2019

Kamvorn W, Techawongstien S, Techawongstien S, Theerakulpisut P. Compatibility of inter-specific crosses between *Capsicum chinense* Jacq. and *Capsicum baccatum* L. at different fertilization stages. Sci Hortic. 2014;179:9–15. https://doi.org/10.1016/j.scienta.2014.09.003.

Kassambara A, Mundt F. factoextra: extract and visualise the results of multivariate data analyses. R package version 1.0.7.; 2020

Katoh K, Standley DM. MAFFT multiple sequence alignment software version 7: improvements in performance and usability. Mol Biol Evol. 2013;30(4):772–780. https://doi.org/10.1093/molbev/mst010.

Khoury CK, Carver D, Barchenger DW, Barboza GE, van Zonneveld M, Jarret R, et al. Modelled distributions and conservation status of the wild relatives of chile peppers (*Capsicum* L.). Divers Distrib. 2020;26(2):209–225. https://doi.org/10.1111/ddi.13008.

Lin T, Zhu G, Zhang J, Xu X, Yu Q, Zheng Z, et al. Genomic analyses provide insights into the history of tomato breeding. Nat Genet. 2014;46(11):1220–1226. https://doi.org/10.1038/ng.3117.

Lin TH, Lin SW, Wang YW, van Zonneveld M, Barchenger DW. Environmental influence on inter- and intraspecific crossability and self-pollination compared to heat treatment of wild and domesticated *Capsicum* species. HortScience 2020;55(9):S109–S110.(abstr).

Loaiza-Figueroa F, Ritland K, Laborde Cancino JA, Tanksley SD. Patterns of genetic variation of the genus *Capsicum* (Solanaceae) in Mexico. Plant Syst Evol. 1989;165(3-4):159–188. https://doi.org/10.1007/BF00936000.

Luna-Ruiz J de J, Nabhan GP, Aguilar-Meléndez A. Shifts in plant chemical defenses of chile pepper (*Capsicum annuum* L.) due to domestication in Mesoamerica. Frontiers Ecol Evol. 2018;6:48 https://doi.org/10.3389/fevo.2018.00048

McCoy JW, Bosland PW. Identification of resistance to powdery mildew in chile pepper. HortScience 2019;54:4–7.

McLeod MJ, Guttman SI, Eshbaugh HW, and Rayle RE. An electrophoretic study of evolution in *Capsicum* (Solanaceae). Evol. 1983;37(3):562–574.

Meyer D, Buchta C. proxy: Distance and similarity measures. R package version 0.4-24.; 2020 https://CRAN.R-project.org/package=proxy

Mongkolporn O, Taylor PJW. Capsicum. In: C. Kole, editor. Wild crop relatives: genomic and breeding resources. Heidelberg, Berlin: Springer; 2011. p. 43–57. https://doi.org/10.1007/978-3-642-20450-0_4.

OECD. Consensus document on the biology of the Capsicum annuum complex (chili peppers, hot peppers and sweet peppers). Paris (France): OECD; 2006

Oyama K, Hernández-Verdugo S, Sánchez C, González-Rodríguez A, Sánchez-Peña P, Garzón-Tiznado JA, et al. Genetic structure of wild and domesticated populations of *Capsicum annuum* (Solanaceae) from northwestern Mexico analyzed by RAPDs. Genet Resour Crop Evol. 2006;53(3):553–562. https://doi.org/10.1007/s10722-004-2363-.

Pickersgill B. Relationships between weedy and cultivated forms in some species of chili peppers (genus *Capsicum*). Evol. 1971;25(4):683–691. https://doi.org/10.2307/2406949.

Pickersgill B. Cytogenetics and evolution of Capsicum L. Chromosome engineering in plants: genetics, breeding, evolution, part B. Amsterdam: Elsevier; 1991

Shipunov A, Murrell P, D’Orazio M, Turner S, Altshuler E, Rau R, et al. shipunov: miscellaneous functions from Alexey Shipunov. R package version 1.12; 2020. https://cran.r-project.org/web/packages/shipunov/index.htm.

Shiragaki K, Yokoi S, Tezuka T. Phylogenetic analysis and molecular diversity of *Capsicum* based on rDNA-ITS region. Horticulturae 2020;6(4):87. doi:10.3390/horticulturae6040087

Tam SM, Lefebvre V, Palloix A, Sage-Palloix AM, Mhiri C, Grandbastien MA. LTR-retrotransposons Tnt1 and T135 markers reveal genetic diversity and evolutionary relationships of domesticated peppers. Theor Appl Genet. 2009;119(6):973–989. https://doi.org/10.1007/s00122-009-1102-6.

Tong N, Bosland PW. *Capsicum tovarii*, a new member of the *Capsicum baccatum* complex. Euphytica. 1999;109(2):71–77. https://doi.org/10.1023/A:1003421217077.

Votava EJ, Nabhan GP, Bosland PW. Genetic diversity and similarity revealed via molecular analysis among and within an in situ population and ex situ accessions of chiltepín (*Capsicum annuum* var. *glabriusculum*). Conserv Genet. 2002;3(2):123–129. https://doi.org/10.1023/A:1015216504565.

Walsh BM, Hoot SB. Phylogenetic relationships of *Capsicum* (Solanaceae) using DNA sequences from two noncoding regions: the chloroplast atpB - rbcL spacer region and nuclear waxy introns. Int J Plant Sci. 2001;162(6):1409–1418.

Wickham H, Chang W, Henry L, Pedersen TL, Takahashi K, Wilke C, et al. ggplot2: Create elegant data visualisations using the grammar of graphics. R package version 3.3.2.; 2020

Yoon BJ, Yang DC, Do JW, Park GP. Overcoming two post-fertilization genetic barriers in interspecific hybridization between *Capsicum annuum* and *C. baccatum* for introgression of anthracnose resistance. Breed Sci., 2006;56(1):31–38. https://doi.org/10.1270/jsbbs.56.31.

van Zonneveld M, Ramirez M, Williams DE, Pretz M, Meckelmann SW, Avila T, et al. Screening genetic resources of *Capsicum* peppers in their primary center of diversity in Bolivia and Peru. PLoS ONE, 2015;10(9). https://doi.org/10.1007/s00217-014-2325-6.

